# Identification of FDA-approved drugs as novel allosteric inhibitors of human executioner caspases

**DOI:** 10.1101/356956

**Authors:** R. N. V. Krishna Deepak, Ahmad Abdullah, Priti Talwar, Hao Fan, Palaniyandi Ravanan

**Affiliations:** Bioinformatics Institute (BII), Agency for Science, Technology and Research (A*STAR), 30 Biopolis Street, #07-01 Matrix, Singapore 138671; Apoptosis and Cell Survival Research Lab, School of Biosciences and Technology, VIT University, Vellore, Tamil Nadu, India, 632014; Department of Biological Sciences, National University of Singapore, 14 Science Drive 4, Singapore 117543; Centre for Computational Biology, DUKE-NUS Medical School, 8 College Road, Singapore 169857

**Keywords:** Apoptosis, Virtual screening, Drug repurposing, Caspase inhibition, NSAIDs

## Abstract

The regulation of apoptosis is a tightly-coordinated process and caspases are its chief regulators. Of special importance are the executioner caspases, caspase-3/7, the activation of which irreversibly sets the cell on the path of death. Dysregulation of apoptosis, particularly an increased rate of cell death lies at the root of numerous human diseases. Although several peptide-based inhibitors targeting the homologous active site region of caspases have been developed, owing to their non-specific activity and poor pharmacological properties their use has largely been restricted. Thus, we sought to identify FDA-approved drugs that could be repurposed as novel allosteric inhibitors of caspase-3/7. In this study, we virtually screened a catalog of FDA-approved drugs targeting an allosteric pocket located at the dimerization interface of caspase-3/7. From among the top-scoring hits we short-listed five compounds for experimental validation. Our enzymatic assays using recombinant caspase-3 suggested that four out of the five drugs effectively inhibited caspase-3 enzymatic activity *in vitro* with IC_50_ values ranging ~10-55 μM. Structural analysis of the docking poses show the four compounds forming specific non-covalent interactions at the allosteric pocket suggesting that these molecules could disrupt the adjacently-located active site. In summary, we report the identification of four novel non-peptide allosteric inhibitors of caspase-3/7 from among FDA-approved drugs.

## INTRODUCTION

Apoptosis or programmed cell death is a vital biological process essential for the elimination of unwanted cells, and maintenance of tissue homeostasis in multicellular organisms. Physiological processes such as embryonic development, normal cell turnover and adult tissue maintenance, immune system function, are closely regulated by apoptosis. Consequently, defects in regulation of apoptosis is commonly associated with various medical conditions such as neurodegeneration, ischemic damage, autoimmune diseases, and certain types of cancers^1,2^. Critical regulators of apoptosis are a family of **c**ysteine-dependent **asp**artic-specific endoprote**ases** called *caspases*. According to their recognized roles in apoptosis, caspases are further sub-classified into initiators (caspase-2, -8 and -9) and executioners (caspase-3, -6, and -7). Among the executioners, caspase-3 is regarded as the primary effector of apoptosis acting as a global mediator essential for multiple proteolytic events^3,4^. Active caspase-3 cleaves an array of protein/peptide substrates triggering apoptosis-associated events such as chromatin condensation and margination, DNA fragmentation, and demolition of structural proteins eventually leading to the death of the cell^5^. Owing to its central role in driving apoptotic pathways, inhibition of caspase-3 has been looked upon as a potential therapeutic strategy to halt progression of a number of diseases including Alzheimer’s disease, Parkinson’s disease, and stroke^6–8^.

Within cells caspase-3 exists as stable but inactive dimeric zymogen or procaspase and is activated via proteolytic cleavage by initiator caspases resulting in a small and a large subunit. Mature caspase-3 molecules are dimers of heterodimers comprising two copies each of the small and large subunits. The substrate-binding region, highly-conserved among caspases, is formed by rearrangement of the active site loops L1-L4 from one heterodimer and L2′ from the other, forming two distinct active sites in each molecule^9^. Several peptide-based caspase inhibitors have been developed which target the substrate-binding/catalytic site (orthosteric)^10^. However, the orthosteric mode of inhibition has the inherent disadvantage of high competition from caspase-3’s diverse indigenous substrates^11^. Most peptide-based inhibitors have also been regarded as mediocre drug candidates for use as pharmacological modulators owing to their poor metabolic stability and cell membrane penetration, thus largely restricting their use to research^12^. On the other hand, exploring an allosteric mode of inhibition presents the possibility of overcoming competitive interference from substrates, and presents with opportunities for the development of small-molecule-based inhibitors with better pharmacological characteristics^13,14^.

The first allosteric site in caspases was discovered by Hardy et al. in caspase-3 employing a site-directed fragment-based ligand discovery approach referred to as disulfide tethering^15^. Using said method two classes of thiol-containing molecules namely, 2-(2,4-dichlorophenoxy)-*N*-(2-mercapto-ethyl)-acetamide (DICA) and 5-fluoro-1*H*-indole-2-carboxylic acid (2-mercapto-ethyl)-amide (FICA) were found to irreversibly inhibit enzymatic activity of caspase-3. Mutational studies, mass spectrometry and peptide mapping studies univocally showed that both DICA and FICA formed a disulfide bridge with a single non-catalytic, surface-exposed cysteine residue (Cys264) located in the central cavity at the dimer interface. Both molecules were also shown to inhibit caspase-7 which shares 53% sequence identity with caspase-3. Structural studies clearly demonstrated the binding of DICA and FICA to the allosteric central cavity *via* covalent interaction with Cys290 in caspase-7 (structurally-equivalent to caspase-3 Cys264) in a 2 ligand:1 dimer ratio. The allosteric site now referred to as exosite A, with bound DICA, along with the active site in capsase-7 are illustrated in Figure 1A. In the crystal structure, two molecules of DICA are symmetrically bound to the allosteric central cavity anchored to the site via specific intermolecular disulfide bridges formed with Cys290 (one from each monomer/chain). Binding of DICA breaks an important cation•••π interaction between Arg187 (large subunit) and Tyr223 (small subunit) which in turn disrupts the organization of the adjacent substrate-binding pocket. Consequently, substrate binding to the active site is affected rendering the enzyme inactive. Structurally, with respect to the conformation of Tyr223 and L2′, the allosterically inhibited caspase structure closely resembles that of the inactive procaspase (Figure 1B)^16^.

**Figure 1.**
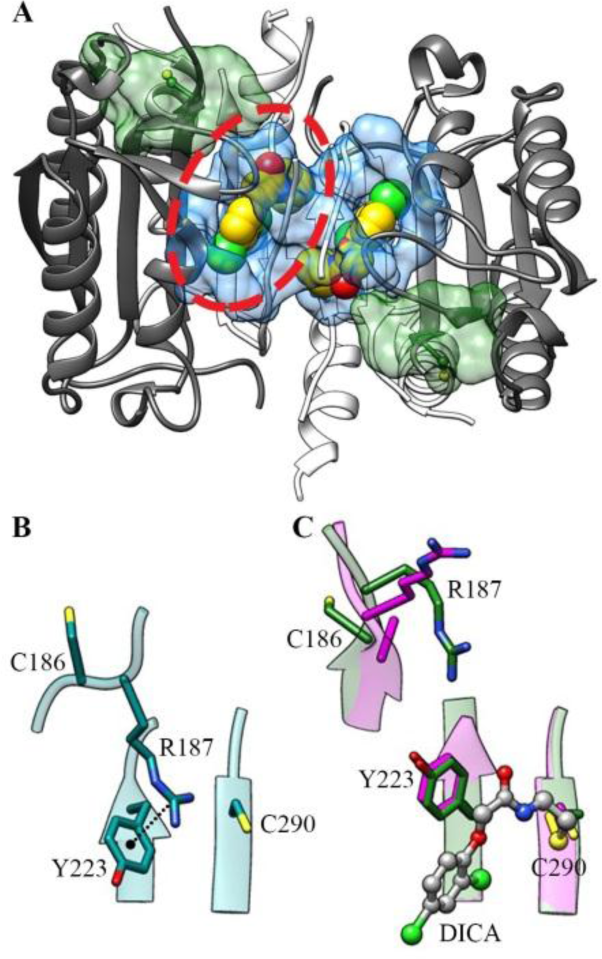
**(A)** Structure of caspase-7 dimer solved in the presence of DICA (yellow). The allosteric central cavity/exosite A and orthosteric active site are depicted using translucent blue and green surface representations, respectively. In our virtual screening, we targeted the region of the allosteric pocket occupied by a single DICA molecule, highlighted with a red dashed line. **(B)** Structure of caspase-7 allosteric site in substrate-mimic DEVD-bound active state (cyan, PDB ID: 1F1J) showing the presence of cation•••π interaction between Arg187 and Tyr223. **(C)** Superposition of structures of caspase-7 allosteric site in DICA-bound inactive state (magenta, PDB ID: 1SHJ), and inactive zymogen state (green, PDB ID: 1GQF).The residue numbers shown correspond to caspase-7 as in PDB ID: 1SHJ for uniformity. Binding of DICA breaks the cation•••π interaction between Arg187 and Tyr223 resulting in a conformation that resembles that of the inactive caspase-7 zymogen.

Although several compounds have exhibited the ability to inhibit caspase-3 activity *in vitro* and in animal models, very few molecules have successfully managed to cross pre-clinical and clinical trials due to their inherent toxicity^17,18^. Therefore, it seems worthwhile to look for potential caspase-3 inhibitors among FDA-approved compounds and understand the mechanism of inhibition with the objective of repurposing them for alternative therapeutic approaches. Thus, we virtually screened a catalog of FDA-approved drugs by docking for their ability to bind to the allosteric site in caspase-7, which is identical to the one found in caspase-3. From among the top hits identified in the virtual screening we selected five compounds for experimental testing their ability to inhibit caspase-3 activity. Four out of the five compounds, namely diflunisal, flubendazole, fenoprofen, and pranoprofen, showed remarkable inhibition of caspase-3 activity *in vitro* both in recombinant caspase-3 activity assay and cell-based assay. In summary, in the present study we demonstrate the value of *in silico* structure-based virtual screening augmented with *in vitro* experimental validation for repurposing FDA-approved drugs as novel allosteric inhibitors for a pharmacologically important but poorly druggable target, namely caspase-3/7.

## MATERIALS AND METHODS

**Preparation of Target Structure for Docking.** No structure of caspase-3 with an allosteric ligand bound in the central cavity is currently available. However, caspase-3 and -7 share high sequence identity (53%) and structural similarity with all 16 residues projecting into the central cavity being identical. Thus, we reasoned that ligands binding to caspase-7 would interact with caspase-3 in a similar fashion. Thus, in this study we targeted the allosteric site in the crystal structure of caspase-7 solved in dimeric form irreversibly complexed with DICA [PDB ID: 1SHJ^15^] in virtual screening. We also screen the same chemical library against the orthosteric site in the caspase-7 structure solved in complex with the peptide inhibitor AC-DEVD-CHO [PDB ID: 1F1J^19^]. Prior to virtual screening, the target structures were prepared by adjusting the side-chain rotamer of Cys290 in chain A of 1SHJ (allosteric site) and Cys186 in chain A of 1F1J (orthosteric site) based on the Dunbrack backbone-dependent rotamer library^20^ breaking the Cys290-DICA and Cys186-AC-DEVD-CHO disulfide bridges, respectively, so that steric clashes between their side-chain sulfur atoms and ligands being sampled could be avoided. Subsequently, all non-protein atoms from the coordinate file the coordinates were removed.

**Virtual Screening.** We screened a catalog of FDA-approved drugs from the ZINC 12 database^21,22^ against the allosteric exosite A in caspase-7 (PDB ID: 1SHJ, chain A) and the orthosteric site (PDB ID: 1F1J, chain A) using DOCK 3.6^23^. We aimed to identify drug molecules that bind preferentially to the allosteric dimerization interface of caspase-3/7 instead of the orthosteric site. For the allosteric site we considered approximately half the molecular surface of exosite A (Figure 1A), formed by residues primarily from chain A of caspase-7, for docking. A total of 45 matching spheres were generated based on the crystallographic coordinates of DICA (PDB ID: 1SHJ, chain A) for the allosteric site and AC-DEVD-CHO (PDB ID: 1F1J, chain A) for the orthosteric site, which were used to guide the docking of ligands to the site.

**Cell Culture and Reagents.** HEK293 cells were obtained from NCCS, Pune, India and cultured in DMEM-containing 10% FBS, 1x antibiotic and antimycotic solution, and 2mM L-glutamine at 37°C and 5% CO_2_ in humid atmosphere. All the test compounds used in this study were procured from Sigma-Aldrich (India). We purchased staurosporine from Alfa Aesar (India) and Z-VAD-FMK from Medchem Express (USA).

**Caspase-3/7 Activity Assay.** For the caspase 3/7 activity assay 10×10^3^ HEK293 cells were seeded on a 96 wells plate (opaque, white) on day 0. Next day, medium was removed from each well and then pre-treated with test compounds for 90 minutes followed by treating the cells with staurosporine. Cells were incubated for 24 h and caspase 3/7 activity assay was performed as per manufacturer’s instruction. Readings were measured using luminometer (Berthold).

**Caspase-3 Enzyme Inhibition Studies.** Recombinant human caspase-3 enzyme was obtained from Enzo Life Sciences (BIOMOL cat. # SE-169) and Caspase-Glo^®^ 3/7 assay reagent from Promega, USA.2U of enzyme was used per well in solid bottom and opaque 96-well-plate containing a reaction mixture formed with 10mM HEPES and 0.1% BSA. Reduction in the luminescence reveals the magnitude of inhibition caused by the test compounds from cleaving Z-DEVD-aminoluciferin by recombinant caspase-3 enzyme. Curve fitting for sigmoidal dose response (variable slope) was done using GraphPad Prism 5.0. IC_50_ values were obtained by interpolating the concentration required for 50% inhibition using dose response curve, in GraphPad Prism 5.0.

**Statistical Analysis.** All data are presented as the standard error of mean (S.E.M) of at least two independent experiments. Statistical comparisons were performed using one-way analysis of variance (ANOVA) followed by Bonferroni’s multiple comparison test (Graphpad Prism, version 5.0) and *p*-value < 0.05 was considered statistically significant.

## RESULTS

**Virtual Screening.** Crystal structures of caspase-7 with DICA and FICA [PDB ID: 1SHJ and 1SHL, respectively]^15^ and Comp-A, a Cu-containing pyridinyl compound [PDB ID: 4FEA]^24^ revealed the existence of an allosteric site in caspases symmetrically occupied by two ligand molecules. We selected the structure of caspase-7 dimer inactivated by the binding of DICA as the target protein in our virtual screening strategy for identification of allosteric inhibitors. We wanted to identify ligands that could bind reversibly to the dimer interface in caspase-3/7 in a 2:1 ratio similar to the binding of DICA, FICA, and Comp-A to caspase-7. Thus, we considered a sub-region of the allosteric central cavity (highlighted in Figure 1A) that surrounds one of the two bound DICA molecules (from chain A) and docked a catalog of FDA-approved drugs obtained from ZINC 12 database^21,22^ to this site using DOCK 3.6^23^. To find compounds with preference to the allosteric site over the orthosteric site, we also docked the FDA catalog to the orthosteric site in the caspase-7 structure solved in the presence of the orthosteric peptide-inhibitor AC-DEVD-CHO [PDB ID: 1F1J]^19^. The docked conformations of the FDA compounds were scored based on an energy function incorporating Poisson-Boltzmann electrostatic, van der Waals, and ligand desolvation penalty terms. Among the top 100 hits for the allosteric site we considered only those compounds that did not appear in the top 100 hits for the orthosteric site. In total, five such compounds were selected from the virtual screening for experimental testing. This ligand candidate selection was done by considering their docking ranks, intermolecular polar interactions, and their chemical similarity with respect to DICA and FICA measured by Tanimoto coefficient (Tc) values. The five compounds are flubendazole, L-tryptophan, fenoprofen, diflunisal, and pranoprofen, all dissimilar from DICA and FICA with Tc values less than 0.4. We also did not consider those FDA-approved compounds that have been previously reported to possess caspase-inhibitory activity in our experimental validation studies. Details of five compounds selected are summarized in Table 1.

**Table 1:**
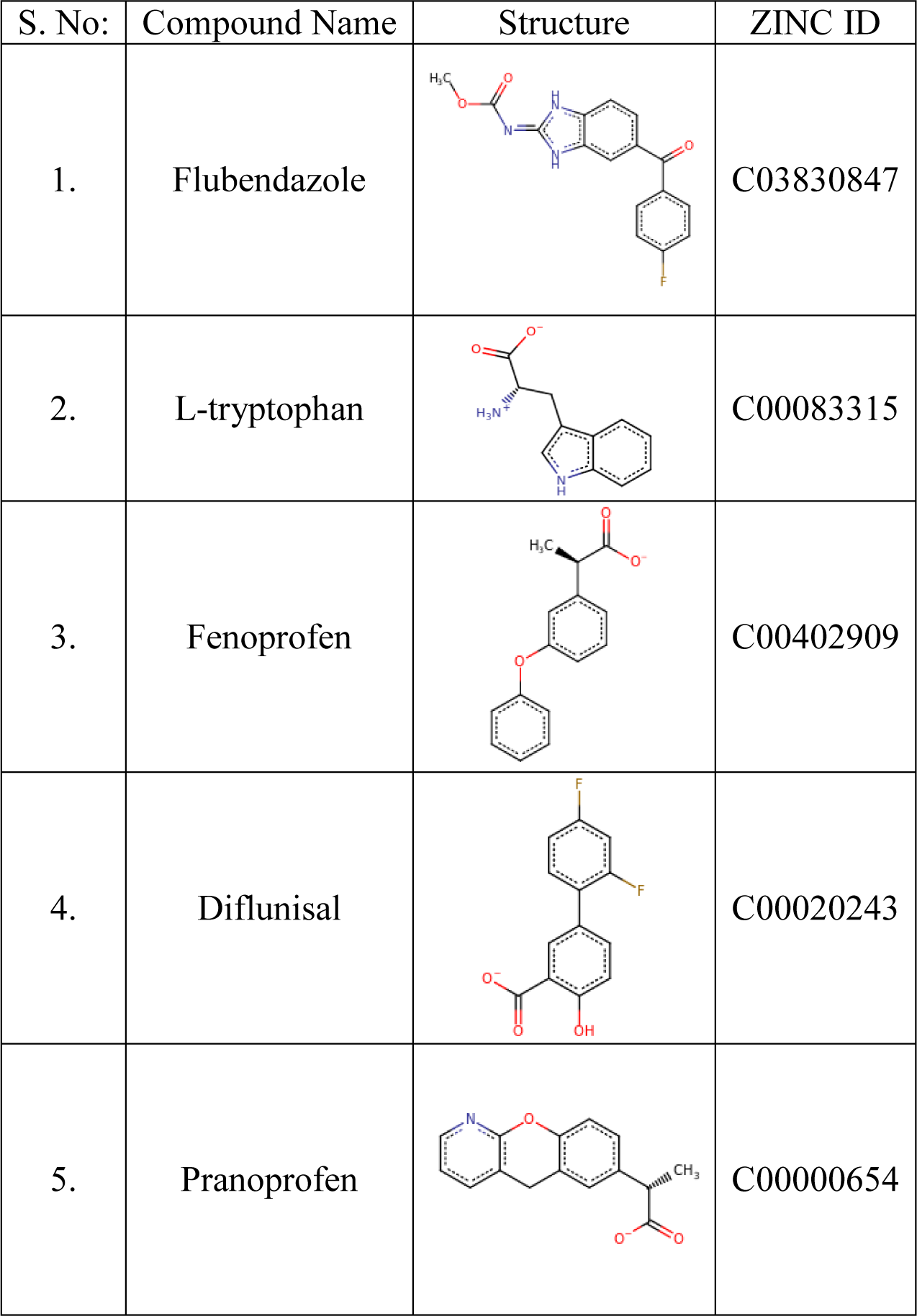
Summary of the five FDA-approved compounds picked for experimental validation of caspase-3 inhibitory activity

| S. No: | Compound Name | Structure | ZINC ID |
| --- | --- | --- | --- |
| 1. | Flubendazole |  | C03830847 |
| 2. | L-tryptophan |  | C00083315 |
| 3. | Fenoprofen |  | C00402909 |
| 4. | Diflunisal |  | C00020243 |
| 5. | Pranoprofen |  | C00000654 |

### Inhibition Studies on Recombinant Human Caspase-3 Enzyme

To test the caspase-3 inhibitory activity of screened hit compounds, Caspase Glo-3/7 assay (Promega) was performed using purified recombinant human caspase-3 enzyme. Since previous studies have ruled out the possibility of NSAIDs inhibiting the activity of luciferase, we did not perform further studies on that prospect^25^. Initial screening was performed using 50 μM concentration of each compound and tested against caspase-3 enzyme activity. Diflunisal and pranoprofen showed maximal inhibition when compared to fenoprofen and flubendazole. L-tryptophan did not show any significant inhibitory activity. AC-DEVD-CHO, a pan-caspase inhibitor, was used as positive control (Figure 2A). The inhibitory activity of the five compounds was studied for up to 60 minutes to estimate their time-dependent inhibitory activity (Figure 2B).

**Figure 2.**
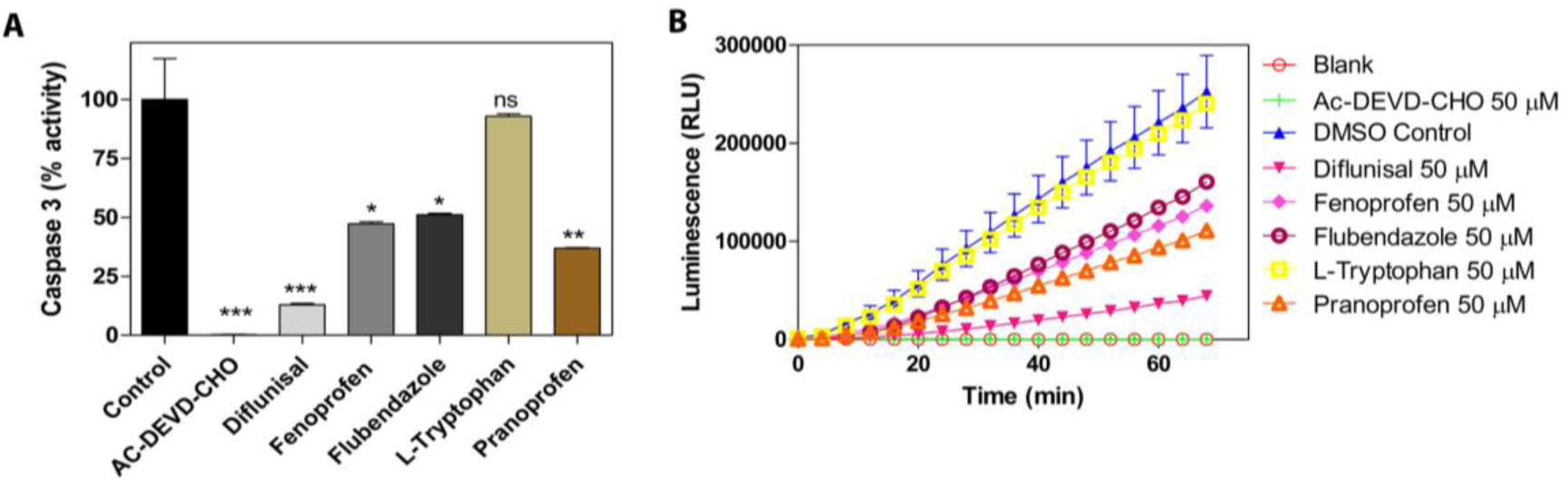
**(A)** Caspase 3/7 activity assay performed with recombinant human caspase-3 enzyme against the five selected compounds for inhibitory activity. **(B)** Time-dependent inhibitory activity of the test compounds. (Ac-DEVD-CHO was used as a positive control). *represents significance between compound added wells compared to control well at p<0.05 (One-way ANOVA).

Further studies on dose-dependent inhibition of compounds also showed a maximum inhibition exhibited by diflunisal compared to pranoprofen, flubendazole, and fenoprofen (Figure 3A-D).

**Figure 3.**
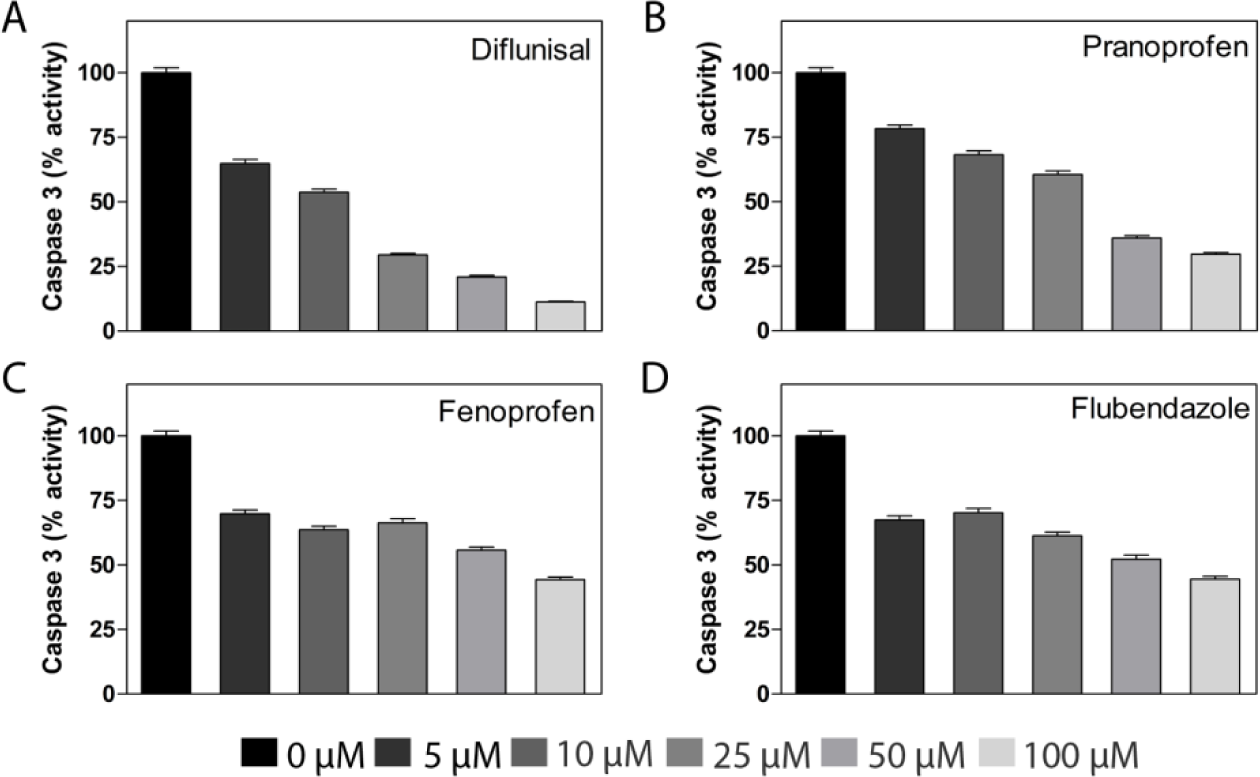
Dose-dependent activity of **(A)** diflunisal, **(B)** pranoprofen, **(C)** fenoprofen, and **(D)** flubendazole showing inhibition of caspase-3 enzymatic activity. Data represented as average S.E.M of experiment performed in triplicate.

To ascertain the IC_50_ values of the inhibitors, compounds were again tested with various concentrations ranging from 1 nM to 500 μM. All the four compounds showed caspase inhibition in time- and concentration-dependent manner (Figure 4). However, we did not observe any decreasing enzymatic activity with increase in the concentration of flubendazole beyond 10 μM concentration. The IC_50_ value of all the compounds have been tabularized in Table 2.

**Figure 4.**
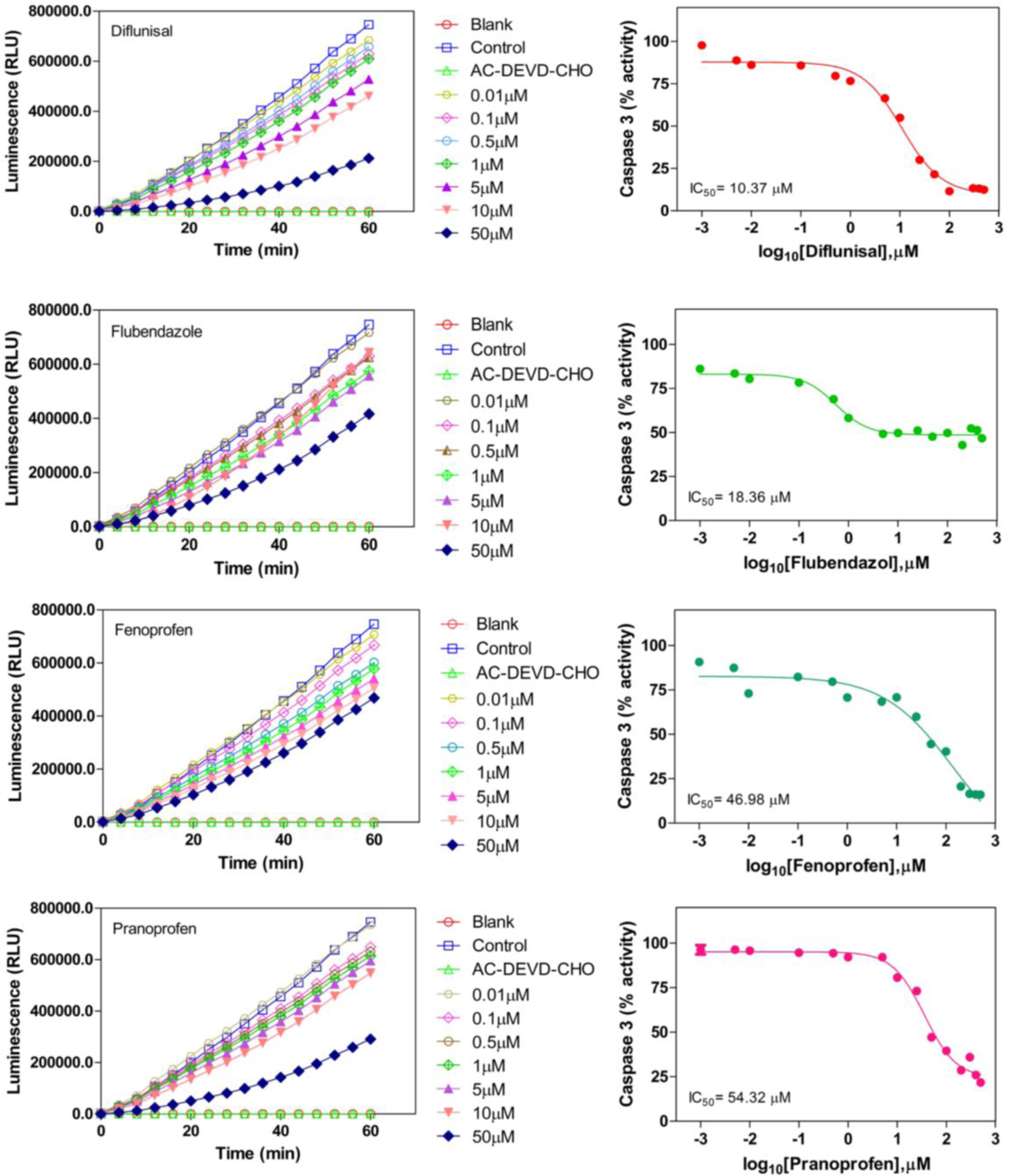
Graphs showing the time- and dose-dependent inhibition of four compounds for a period of 60 minutes and IC_50_ value determination by dose-response curves.

**Table 2:** IC_50_ values determined for the four hit compounds for caspase-3 inhibition activity

| S. No: | Compound Name | Inhibition of caspase-3 |  |
| --- | --- | --- | --- |
|  |  | IC<sub>50</sub> (μM) | R<sup>2</sup> |
| 1. | Diflunisal | 10.37 ± 0.5 | 0.9845 |
| 2. | Flubendazole | 18.36 ± 4.23 | 0.9654 |
| 3. | Fenoprofen | 46.98 ± 6.07 | 0.9657 |
| 4. | Pranoprofen | 54.32 ± 0.6 | 0.9838 |

From Table 2 it is clear that diflunisal exhibits the best inhibition profile against caspase-3 *in vitro* followed by flubendazole, fenoprofen and pranoprofen. In order to speculate on the structural basis of the inhibitory capacity of the tested compounds, we analyzed the various non-covalent interactions formed between the docked compounds and the residues of the allosteric pocket. The docking poses of the four compounds showing caspase-3 inhibitory activity are illustrated in Figure 5. From the figure it is evident that all four compounds form extensive intermolecular non-covalent interactions within the allosteric binding site.

**Figure 5.**
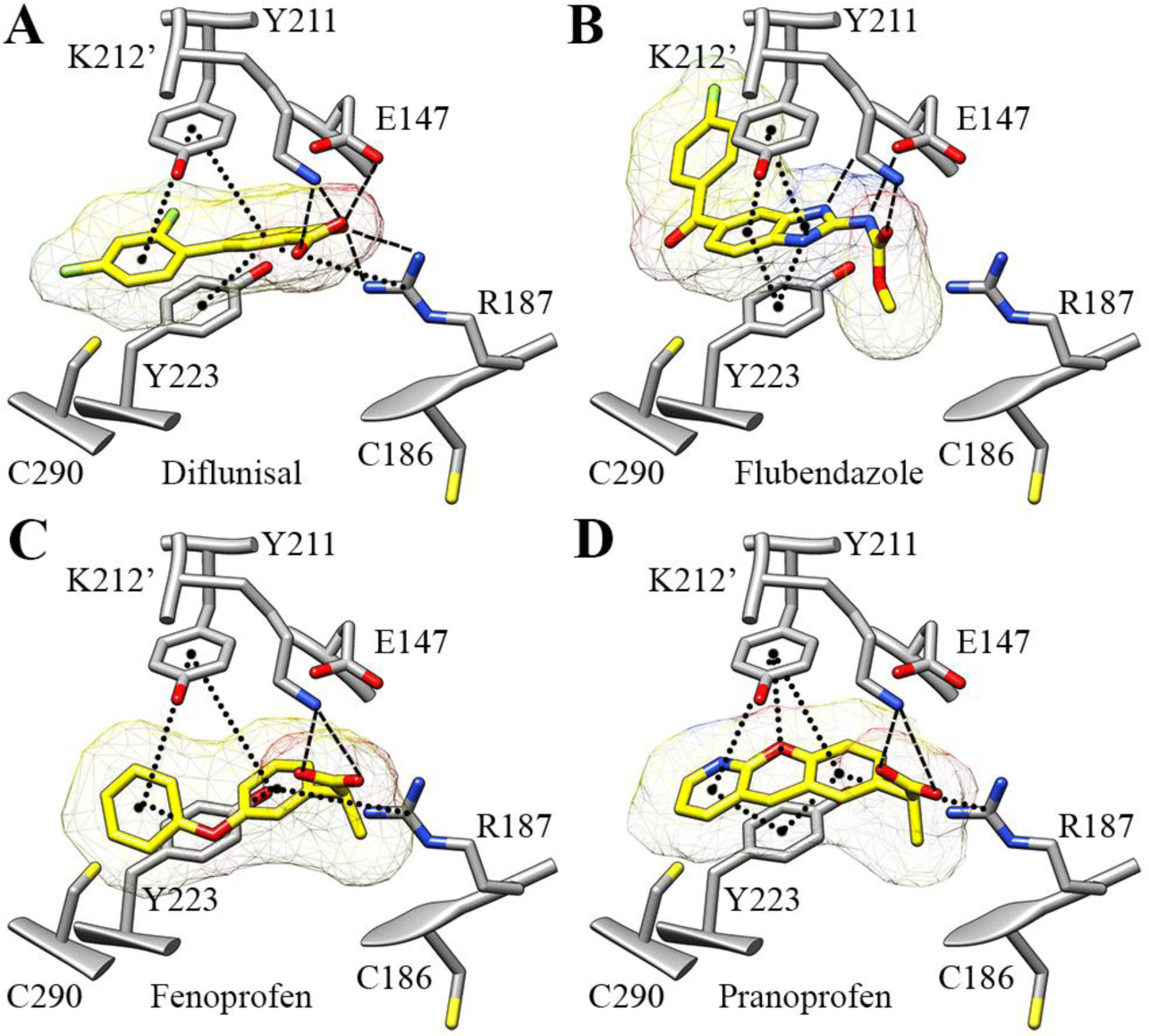
Docking poses of the four compounds showing caspase-3 inhibitory activity, namely (**A)** diflunisal, (**B)** flubendazole, (**C)** fenoprofen, and (**D)** pranoprofen within the allosteric pocket in caspase-7 (PDB ID: 1SHJ, chain A). Non-covalent interactions such as π•••π stacking and cation•••π are shown using dotted lines while salt-bridges and hydrogen bonds are shown using dashed lines. The geometric criteria for identifying the interactions have been used from Jain et al 2011. The active site C186 and allosteric site C290 are shown for reference.

Our analysis revealed the presence of various intermolecular interactions such as hydrophobic contacts, π•••π, cation•••π, salt-bridge, and hydrogen bonds in the docked complexes (Figure 5). Overall the allosteric pocket in each monomer can be considered as having an interior hydrophobic and an exterior polar region. The hydrophobic region comprises Tyr211, Ile213, Tyr223, Cys290, Val292, Val215′, and Val292′, whereas the polar region includes Glu146, Glu147, Arg187 and Lys212′. All four compounds, in their docked conformations, are aligned such that one of their aromatic moieties is buried in the hydrophobic region of the allosteric pocket while their polar moieties are favorably positioned towards the exterior polar region. Based on this observation it could be speculated that L-tryptophan did not show any inhibitory activity owing to its smaller size preventing it from fully occupying the allosteric pocket.

All four compounds showing inhibitory activity have two aromatic moieties both of which form π-stacking interactions with Tyr211 and Tyr223. Except flubendazole, docked conformations of the three other compounds shows the presence of a cation•••π interaction with Arg187 suggesting that an ability to engage Arg187 could be crucial for effecting capase-3 inhibition. Although the docked conformation of flubendazole doesn’t form the cation•••π interaction with Arg187, it forms multiple H-bonds with Glu147, which in turn could stabilize its interaction with caspase-3. It could thus be speculated that binding of the inhibitory compounds could break the intramolecular cation•••π interaction between Arg187 and Tyr223 observed in active caspase-3/7 (Figure 1B) and engage the residues in intermolecular interactions (Figure 5), thereby disrupting the organization of the active site similar to DICA (Figure 1C). The binding of the four compounds could also be further favored by the formation of salt-bridge/hydrogen bonds with residues such as Glu146, Glu147, and Lys212′. In summary, it could be suggested that small-molecules that possess the features to disrupt the Arg187•••Tyr223 interaction and engage these residues in favorable interactions (*via* aromatic moieties) along with Glu147 (*via* hydrogen bond donors), and Lys212′ (*via* carboxylate group) could act as allosteric inhibitors of caspace-3/7.

### Analysis of Hit Compounds for Caspase-3/7 Inhibition Activity in HEK293 cell line

To confirm the caspase-3 inhibitory activity of the four compounds *in* vitro, we assayed the compounds against HEK293 cells treated with staurosporine. Staurosporine is a pan-kinase inhibitor and a well-known activator of caspase-3 activity. Our results showed that while all four compounds exhibited inhibition of staurosporine-induced caspase-3 activity in HEK293 cells, diflunisal at 50 μM concentration exhibited maximal inhibition (Figure 6).

**Figure 6.**
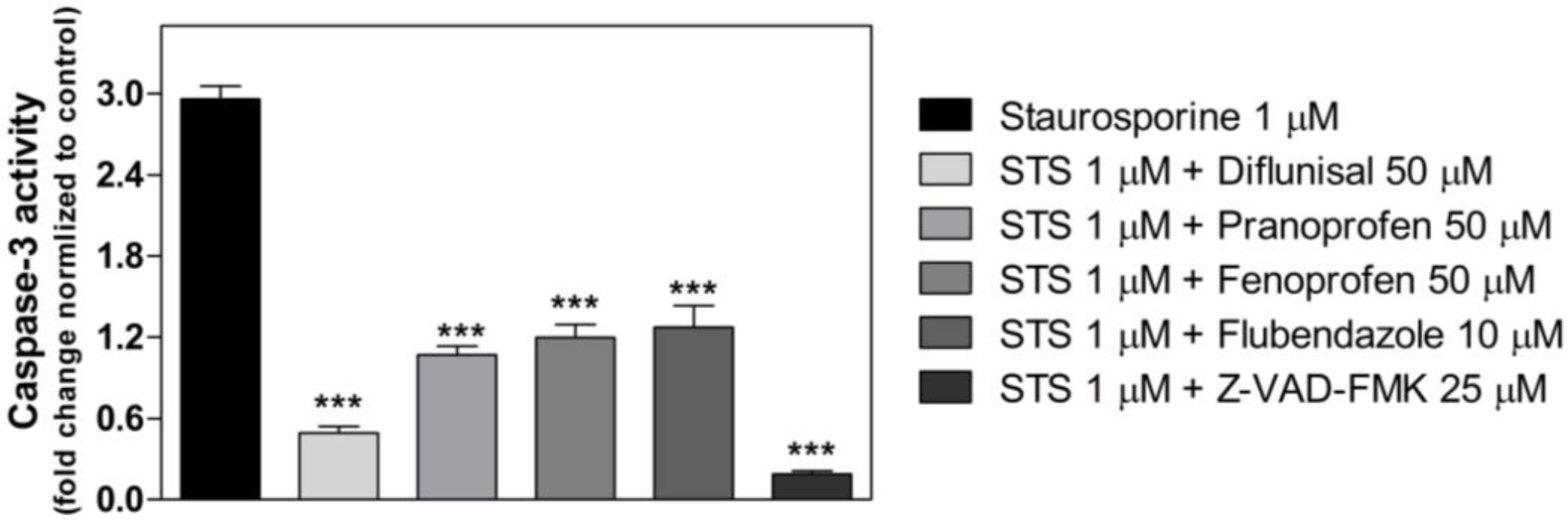
Caspase 3/7 activity assay showing the fold increase of caspase 3/7 activity in Staurosporine (STS 1 μM) alone treated HEK293 cells and inhibition showed by pretreatment with screened compounds after 24h of incubation. * represents the significance between staurosporine alone treated cells to compounds pretreated cells at p< 0.05 (One way ANOVA).

## DISCUSSION

Structure-guided virtual screening is a well-established approach and has been used successfully in numerous earlier studies for the identification of allosteric modulators of important pharmacological targets^26,27^. In the present study, we prospected a catalog of FDA-approved drugs using a structure-based virtual screening approach and identified novel reversible allosteric inhibitors targeting the human caspase-3/7. We specifically targeted the allosteric exosite A on the dimerization interface of caspase-3/7 in order to identify compounds that could bring about inhibition of enzyme function. *In vitro* functional assays of capsase-3 enzyme activity showed that four out of the five short-listed FDA-approved compounds, namely diflunisal, pranoprofen, flubenazole, and fenoprofen exhibited inhibitory capabilities, with diflunisal and flubendazole showing remarkable potency. The common features among these compounds are the presence of two or more aromatic rings, a carboxylate group, and hydrogen bond donors. Based on the docking poses observed for the compounds, it could be suggested that intermolecular π•••π stacking interactions with Tyr211 and Tyr223 and salt-bridge/polar interactions with Glu147, Arg187 and Lys212′ of exosite A could favor ligand binding. The essential role of π•••π stacking, cation•••π interactions and other non-covalent interactions in ligand binding has been extensively studied in numerous protein-ligand systems ^28-32^.

Since the reports on DICA and FICA the dimerization sites of executioner caspases have been targeted for designing non-peptide caspase inhibitors, but no small-molecules have been approved for use as drugs against caspases yet^33^. Unlike DICA and FICA, which binds irreversibly to caspase-3/7, compound-A reported by Feldman et al interacts reversibly with the dimerization site of executioner caspase-7^34^. Recently, Spies and co-workers^35^, using a fragment-based drug discovery approach, identified a series of compounds exhibiting allosteric inhibitory capabilities against caspase-7 (IC_50_ values 637-8520 μM). Two of these compounds share good structural similarity with three compounds tested in our study (Tc value 0.4-0.7) and are found to occupy part of the allosteric pocket at the dimer interface in caspase-7. Results from the study strongly support our conclusion that the compounds identified in our study could indeed bind to the dimer interface of caspase-3/7 and bring about reversible allosteric inhibition suggesting that these FDA-approved drugs can be taken further as therapeutic leads. It is also interesting to note that three out of the four FDA compounds capable of inhibiting caspase-3 activity, namely diflunisal, pranoprofen, and fenoprofen, are non-steroidal anti-inflammatory drugs (NSAIDs). We also observed other NSAIDs among the top 100 hits from our virtual screening, such as fenbufen, ibuprofen, and flufenamic acid which have been shown to have anti-caspase activity by Smith et al^36^. To our knowledge this is the first study to reporting the ability of FDA-approved drugs to act as allosteric inhibitors of caspase-3 targeting exosite A. The combined *in silico-in vitro* approach presented here could help circumvent the limitations of traditional peptide-based inhibitors such as poor pharmacological properties and metabolic stability. The pharmacophores of the newly-identified inhibitors could also help in the identification/design of specific and more potent caspase inhibitors. For example, a search of the ZINC 12 database for compounds with 70% structural similarity to diflunisal lists 176 compounds that are commercially available. Screening of these compounds could bring to fore new molecules that have better potency and selectivity. To summarize, our study reports the identification of four FDA-approved drugs that can allosterically inhibit caspase-3/7 and establishes a new direction for researchers to explore for rational drug design and development.

## Author Contributions

P.R., H.F., P.T., A.A. and K.D. conceived and designed the experiments. K.D. and A.A. performed the computational and experimental studies, respectively, and are co-first-authors contributing equally to the work. A.A., K.D., P.R., and H.F. analyzed and interpreted the data, and wrote the manuscript.

## Notes

The authors declare no competing financial interest.

## ACKNOWLEDGEMENTS

K.D. and H.F. thank Dr. Fu Lin of Fan lab for his help in hit-picking. K.D. and H.F. greatly acknowledges the financial support from Biomedical Research Council of A*STAR, Singapore. The computational work for this article was partially performed on resources of the National Supercomputing Centre, Singapore (https://www.nscc.sg). P.R. greatly acknowledges the financial support from Science and Engineering Research Board (SB/EMEQ-223/2014), Department of Science and Technology, India.

## ABBREVIATIONS USED

DICA, 2-(2,4-Dichlorophenoxy)-*N*-(2-mercapto-ethyl)-acetamide; FICA, 5-Fluoro-1*H*-indole-2-carboxylic acid (2-mercapto-ethyl)-amide; NSAID, non-steroidal anti-inflammatory drugs.

## SUPPORTING INFORMATION

Supporting information available: Multiple sequence alignment of caspases.

